# Inferring Bacterial Interspecific Interactions from Microcolony Growth Expansion

**DOI:** 10.1101/2024.05.19.594856

**Authors:** Tania Miguel Trabajo, Isaline Guex, Manupriyam Dubey, Elvire Sarton-Lohéac, Helena Todorov, Xavier Richard, Christian Mazza, Jan Roelof van der Meer

## Abstract

Interactions between species are thought to be crucial for modulating their growth and behaviour within communities, and determinant for the emergence of community functions. Several different interaction concepts exist, but there is no consensus on how interactions should be quantified and integrated in community growth theory. Here we expand on existing concepts of real-time measurements of pure culture microcolony growth to develop and benchmark coculture microcolony experiments, and show how these can both parametrize growth kinetic and interspecific interaction effects. We follow surface growth by time-lapse microscopy of fluorescently tagged *Pseudomonas putida* and *Pseudomonas veronii* under substrate competition with succinate, or under substrate indifference with D-mannitol and putrescine. Monoculture-grown microcolonies showed substrate concentration dependent expansion rates as expected from Monod relations, whereas individual microcolony yields were strongly dependent on densities and spatial positioning of founder cells. Maximum specific growth rates in cocultures under substrate competition were diminished by ca. 15%, which was seeding-density independent. The collective *P. putida* population dominated growth over that of *P. veronii*, but with 27% yield loss under competition compared to monoculture growth; and 90% for that of *P. veronii*. Incidental local reversal of competition was observed where *P. veronii* microcolonies profited at the detriment of *P. putida*, and between 9 and 43% of *P. veronii* microcolonies grew bigger than expected from bulk competition, depending on seeding density. Simulations with a cell-agent Monod surface growth model suggested that colony expansion rate decrease in competitive coculture is caused by metabolite cross-feeding, which was supported by exometabolite analysis during and after growth of the strains on their individual or swapped supernatant. Coculture microcolony growth experiments thus provide a flexible platform for analysis of kinetic and interspecific interactions, expanding from individual microcolony phenotypic effects to averaged behaviour across all microcolony pairs. The system in theory is scalable to follow real-time growth of multiple species simultaneously into communities.

## INTRODUCTION

Understanding how interactions between microbial cells of different species shape the formation and functioning of multi-species communities, remains a long coveted goal in microbial ecology (Faust & Raes, 2012, Coyte *et al*., 2015, Zelezniak *et al*., 2015, Kehe *et al*., 2021, Schäfer *et al*., 2023). Currently, however, there is no consensus in the field on how interspecific interactions should be described or quantified (Pacheco *et al*., 2022). Depending on the data type and the theoretical framework, interaction concepts have been based on, for example, species co-occurrence networks (Faust & Raes, 2012, Widder *et al*., 2022), machine-learned community compositional patterns (Emmenegger *et al*., 2023), resource utilization (Piccardi *et al*., 2019, Rodríguez Amor & Dal Bello, 2019, Dal Bello *et al*., 2021, Nestor *et al*., 2023) and nutrient niche overlap predictions (Schäfer *et al*., 2023), metabolite exchange (Zelezniak *et al*., 2015, Pacheco *et al*., 2019, Kehe *et al*., 2021), or growth expansion patterns (Momeni *et al*., 2013, Goldschmidt *et al*., 2017, Borer *et al*., 2020), and even on cell-cell contact secretion systems (Basler *et al*., 2013, Niggli *et al*., 2021). One of the difficulties to extrapolate from different interaction levels to community formation, is their integration across spatial scales and time, and their embedment in appropriate growth kinetic frameworks (van den Berg *et al*., 2022). Connecting here to classical Monod-type growth kinetics would make sense, given its prevalence and acceptance in the context of pure culture physiology, and recent adaptations to the level of multispecies growth (Goldford *et al*., 2018, Liao *et al*., 2020, Dal Bello *et al*., 2021, van den Berg *et al*., 2022, Guex *et al*., 2023). However, co-or multispecies culture data adaptable to multispecies Monod-type growth kinetic frameworks are still scarce, as one needs appropriate parameter values for species-individual growth rates and yields, their dependencies on nutrient conditions, and their variability under influence of emerging interspecific interactions. To achieve this, we propose here to adapt an existing, scalable methodology to extract kinetic parameters from microcolony growth patterns, similar as have been used in pure culture studies in the area of environmental, food and medical microbiology (Reinhard & van der Meer, 2010, Eijlander & Kuipers, 2013, Koutsoumanis & Lianou, 2013, Nghe *et al*., 2013, Jung & Lee, 2016).

Whereas growth kinetic measurements for two or more species in mixed liquid cultures are rather cumbersome, because of the difficulty to identify the individual species’ contributions, real-time microcolony growth measurements are much easier, because the species can be differentiated by their spatial position, while the dividing cells can remain sufficiently close to allow interactions to emerge. Quantifying growth and interactions at microcolony level would have the additional advantage of maintaining similar spatial scales that most bacterial cells face in natural, heterogeneous habitats and under resource-diffusion limited growth conditions. In the context of unsaturated soil, for example, microbial growth habitats are characterized by small water-filled pockets, pores and water-film covered surfaces (Tecon & Or, 2017). It has been estimated that the majority of cell clusters in unsaturated soils have fewer than 100 cells (Bickel & Or, 2023), with inter-cell distances averaging between 10 and 100 µm (Raynaud & Nunan, 2014). Interspecific interactions are, therefore, also expected to emerge across short distances, with cells, potentially, going only through short growth cycles as a result of limited carbon, water and space.

The objectives of this research were to expand a microcolony growth framework from single pure to mixed cultures, such that relevant kinetic parameters can be extracted of the individual species in the mixture and emerging interspecific interactions can be quantified across relevant local scales (10–100 µm inter-cell distances). Our set-up consists of microscopic growth chambers with cells growing on miniature nutrient surfaces (Reinhard & van der Meer, 2010, Reinhard & van der Meer, 2014) that can be operated under long incubation times (2-4 days), enabling real-time microscopy of all growth phases from individual founder cells to mature microcolonies (i.e., lag times, exponential growth, and stationary phase). To benchmark the concept, we deployed two fluorescently labelled soil bacteria: *Pseudomonas veronii* (Morales *et al*., 2016) and *Pseudomonas putida* (Zylstra & Gibson, 1989), and quantified growth in individual mono-and cocultures under both competitive (i.e., same primary substrate or shared metabolites) or substrate indifference conditions (i.e., each species has its own unique substrate). In addition, we developed an individual cell-agent model based on Monod growth under substrate-diffusion conditions and with inclusion of interspecific interactions (Guex *et al*., 2023), that we parametrised using the empirical kinetic data in order to understand growth and interaction effects both at the scale of local individual microcolonies, and at the level of the community as a whole (the ensemble of all microcolonies on the growth surface). Our results indicate good agreement of averaged microcolony growth in mono- and cocultures and bulk population competition measurements, but surprising effects of variations in single (founder) cell kinetic parameters on the local reproductive success. Contrary to expectations from Monod theory, we find that competitive interactions not only reduce microcolony yields of either partner, but also their maximum specific growth rates. Although shown here for a defined coculture, the same microscopic growth platform is scalable to more complex mixed cultures.

## MATERIALS AND METHODS

### Study system design for interspecific interaction measurements from microcolony growth

We deployed a closed sterile microscopy chamber with a 1-mm thin agarose-solidified disk (‘agarose patch’, Fig. 1A), containing low carbon substrate concentrations (1 mM, to avoid multi-layered growth), onto which individual cells are randomly and sparsely seeded (Reinhard & van der Meer, 2014). Cell division into microcolonies is imaged and recorded by time-lapse epifluorescence microscopy (Fig. 1A), from which colony expansion rates, lag times, final colony size area, colony distances and other relevant parameters are extracted and compared between individual mono- and co-culture incubations. The system was benchmarked with *P. putida* (Ppu) and *P. veronii* (Pve), cultured either under conditions of (assumed) direct substrate competition or substrate indifference. To induce substrate competition, we added succinate to the agarose patch, whereas substrate indifference was generated by adding both D-mannitol, as a specific substrate for Pve, and putrescine for Ppu. Local biomass formation (by microscopy) for either strain alone or in coculture was compared to bulk yields across the whole patch, by washing cells from the surfaces at the end of the experiment and counting either species by flow cytometry on the basis of its fluorescent marker.

**Figure 1.**
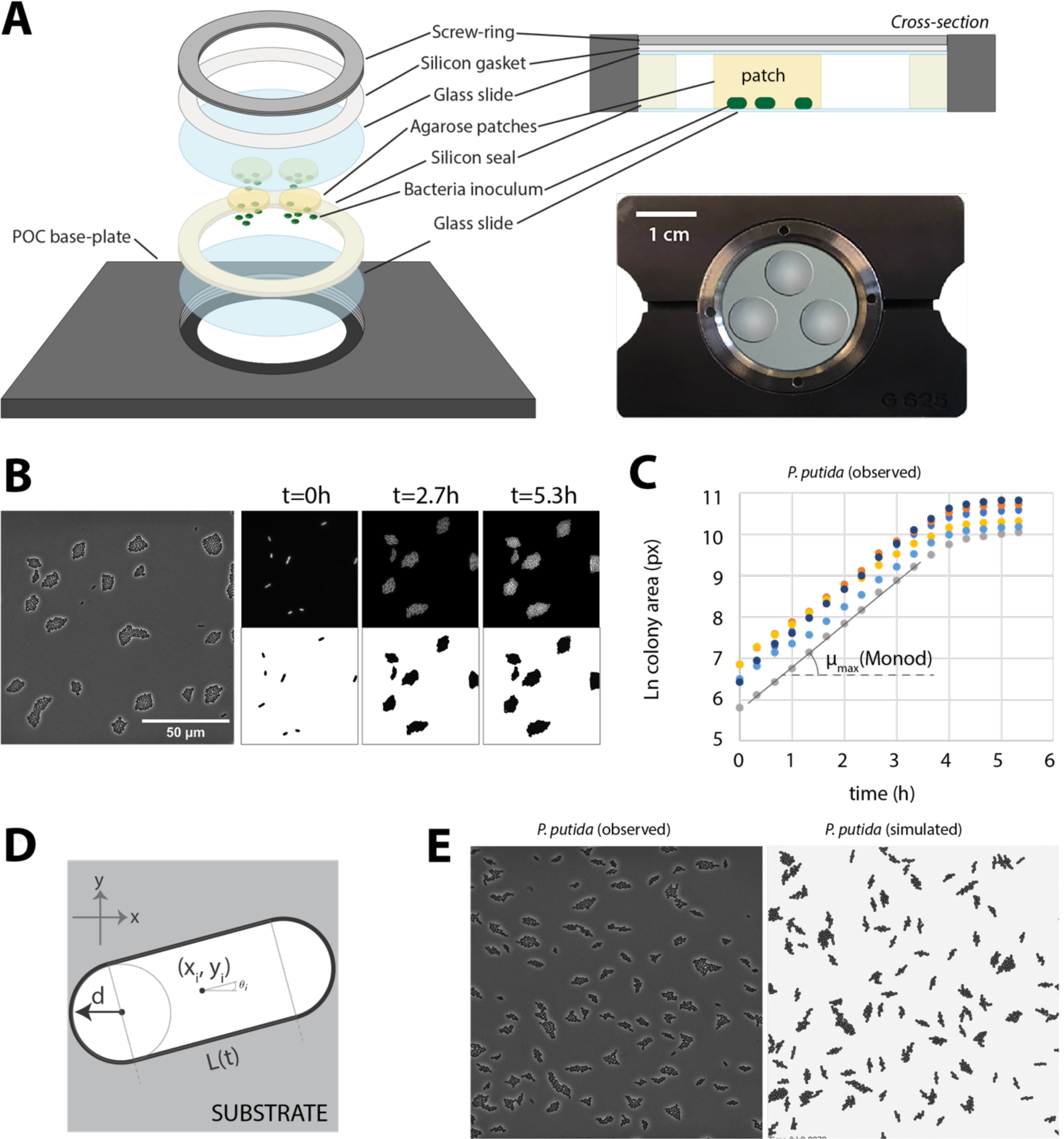
Set-up and principle of the microcolony growth expansion experiments. A) Closed microcolony growth chamber to incubate the agarose patches and the surface-deposited cells for real-time microscopy. B) Example of an imaged surface area with *P. putida* microcolonies; snapshots taken at difference time points and the resulting colony segmentation. C) Deduction of maximum specific growth rates from ln-transformed segmented microcolony areas over time. D) Cell-agent used for the Monod-based growth model, assuming cylindrical cells expanding in x,y direction on the surface. E) Comparison of observed and simulated *P. putida* microcolonies at the same seeding positions and incubation times.

### Bacterial strains and preparation of founder cell cultures

*P. putida* F1 (Zylstra & Gibson, 1989), was genetically labelled with a single copy mini-Tn*5* insertion (Martinez-Garcia *et al*., 2011) constitutively expressing eGFP under the control of the ICE*clc*-promoter P_circ_ (Sentchilo *et al*., 2003) and containing a gene for kanamycin resistance. *P. veronii* 1YdBTEX2 (Morales *et al*., 2016) was labelled with a single copy mini-Tn*7* insertion constitutively expressing mCherry under the control of the P_tac_ promoter and providing gentamicin resistance (Rochat *et al*., 2010). Cultures were stocked at –80 °C in 15% *v/v* glycerol and regrown for each experiment on Nutrient Agar medium (Oxoid CM 0067) with the appropriate antibiotics to obtain individual colonies. A single fresh colony was then precultured in liquid 21C minimal medium without vitamins (MM) (Gerhardt *et al*., 1981) supplemented with 5 mM of the carbon source to be tested in the patch, with the appropriate antibiotic included, and incubated at 30 °C with rotary shaking (160 rpm). Cells were harvested from a 2 ml-aliquot sampled from exponentially growing (culture density of OD_600_ = 0.8) or from stationary phase cultures. The aliquot was centrifuged for 2 min at 13,000 rpm in a Heraeus Fresco 21 microfuge (Thermo Scientific) at room temperature. The supernatant was decanted, and the cell pellet was resuspended in 1 ml MM (without carbon source). This procedure was repeated twice more. After the final resuspension, the culture turbidity was again measured, and suspensions were diluted with MM to an OD_600_ of 0.07 for Ppu and 0.11 for Pve. Cell numbers were quantified further by flow cytometry (see below). These suspensions were then used directly for monoculture patch seeding or mixed in a 1:1 *v/v* ratio to produce cocultures with approximately equal founder cell numbers.

### Preparation of microcolony growth surfaces

Molten agarose solution was prepared with 10 g l^−1^ UltraPure™ Agarose (Invitrogen, 16500-100) in MM, to which the desired carbon source was added while the agarose mixture was still liquid at 45 °C. Stocks of individual carbon sources (succinate, 490 mM; putrescine, 33 mM and D-mannitol, 150 mM) were prepared by weighing from the pure substance (Sigma Aldrich) in ultrapure water, which was sterilized by passing through a 0.2-µm membrane filter (ClearLine, 037044). Carbon substrates were diluted in the agarose solution to achieve final concentrations of: 0.01 mM, 0.05 mM, 0.5 mM, 1 mM or 5 mM (for succinate); or 0.66 mM for putrescine plus 1 mM for D-mannitol (i.e., to have equal C-molarity for the indifference scenario). Agarose without any added carbon substrate served as a control for background growth.

A volume of 1.5 ml of agarose-carbon substrate solution at 45 °C was poured on a circular 42– mm diameter and 0.37–mm thick glass slide (H. Saur Laborbedarf, Germany) enclosed with a 1-mm thick silicon ring (Fig. 1A) (Reinhard & van der Meer, 2010). Immediately after pouring, a second glass slide was placed on top to give the patch the desired thickness. After 5 min solidification, the top slide was gently removed and multiple 1-cm ø circular disks were punched from the solid agarose using a steel rod cutter previously sterilized with the flame. Circular disks were arranged on a new round glass slide (max. 4 per slide, Fig. 1A). An aliquot of 10 µl of diluted founder cell suspension (either mono-or coculture mixture), was then carefully pipetted on each surface and spread homogenously, and dried for 10 min in a laminar sterile flow hood at 21 °C. The slide with patches was turned upside down on a new clean round glass coverslip with a separation silicon ring of 1 mm thickness, which was mounted in a sterilized black anodized chamber (POC chamber; H. Saur Laborbedarf, Germany). A further 0.5-mm-thick silicon gasket was placed on the top glass coverslip, before closing the chamber with a screw ring (without putting pressure on the slide). In the final setup (Fig. 1A), cells are caught in between the lower cover slip and the agarose surface, whereas the top of the agarose touches the upper coverslip (i.e., no air space in between to avoid condensation droplets; Fig. 1A). The remainder of the chamber has ambient air that diffuses into the patches from the sides.

### Time-lapse imaging of microcolony growth

Microscope chambers were mounted (cells facing down) and incubated on a Nikon ECLIPSE Ti Series inverted microscope coupled with a Hamamatsu C11440 22CU camera and a Nikon CFI Plan Apo Lambda 100× Oil objective (1.45 numerical aperture, 1000× final magnification). The temperature was kept between 22-24 °C. Time-lapse programming was controlled by a script in MicroManager Studio version 1.4.23. Images in phase contrast, GFP or mCherry fluorescence were taken every 20 min, with exposure times of 30 ms, 50 ms and 100 ms, respectively, and using a 4% power-set pE-100 LED illumination system from CoolLED. Imaging positions (between 8 and 10 per patch) were defined randomly at the start, but within a 3-mm radius of the patch centre to avoid edge effects, and then kept for the remainder of the experiment. The total duration of the growth varied between experiments, from 12 h (in case of imaging only exponential growth) to 72 h (capturing stationary phase microcolonies). Images were saved as 16-bit .tif files for each channel, per position and per time point.

### Image analysis and extraction of growth kinetic parameters

Microcolony growth was extracted from the fluorescent time-lapse image files, which were processed in a custom-made Python script (version 3.8.3) through a Jupyter Notebook (version 6.0.3). Colonies were segmented at each time step on the fluorescent images using the Otsu algorithm with a variable threshold that adapts to every frame. Segmented areas were stored as individual objects (in pixels), aligned across images, and then overlaid from a complete time-series in order to extract colony area expansion rates (Fig. 1B, C). Non-dividing single cells (objects with the same size at the beginning and at the end of the time-lapse) were removed by filtering, but single elongating cells were included (even if eventually they did not divide). Microcolonies positioned near the image edges (within 200 px from the border) were removed to avoid taking incomplete microcolonies into account.

Maximum colony expansion rates (*r*) were calculated from the exponential growth phase as the mean of moving linear slopes of ln-transformed colony areas versus time, including at least 5 consecutive time points and only slopes with an r^2^ >0.99. Colonies with starting areas <150 px (which can be the result of faulty image overlays) were corrected to the minimum cell size (=150 px) before calculating ln-transformed slopes. Since colony area is a proxy for cell biomass, the maximum colony expansion rate *r* from the ln-transformed areas was taken as the *µ_max_* of biomass growth of the respective colony.

### Flow cytometry quantification of total cell numbers

To quantify the final (combined) cell numbers of either strain growing on the agarose patches, the cells were washed from the surface at the end of the experiment by dismounting the POC chamber, flooding each individual patch with 2 ml MM, and pipetting repeatedly to resuspend the cells. This suspension was subsequently serially diluted in MM and aspired on a NovoCyte Flow cytometer (OMNI Life Science Agilent) at a flow rate of 14 µl min^−1^ and a total analysed volume of 15 µl. Strains were gated based on their specific fluorescence using the NovoExpress software (v. 1.4.1). GFP was measured with the instrument’s FITC channel (excitation at 488 nm and emission/detection at 530 nm), whereas mCherry fluorescence was detected in the PE-Texas Red channel (561 and 615 nm).

### Individual cell-agent surface growth model

To better understand the underlaying factors causing growth kinetic variations in mono and cocultures, we simulated microcolony growth on the agarose patch in an agent-based surface growth kinetic model (Fig. 1D, E and Appendix A). Individual cell-agents (with volumes similar to Ppu or Pve) are positioned on a grid surface with similar dimensions as the experimentally imaged surface areas (134 × 134 µm^2^). Users can chose to set the founder cell densities in the model following the experimentally observed geometric cell centre positions, or at random positions. Cells are given inherent growth kinetic properties (i.e., *µ_max_*, K_s_ and yield) by sampling from experimental observations or by *a priori* definition, allowing cell-to-cell variations and/or individual lag times (i.e., before the onset of cell growth). The model then assumes substrate to biomass conversion following Monod growth and yield, which is translated into cell elongation and division. Simulations calculate changes in resources, metabolites, cell biomass and positions for each time step *Δt* (corresponding to ca. 0.72 s) and per volumetric box of 3.7 pL in the surface grid. Molecular diffusion across boxes is recalculated after every time step. Cells reposition after each time interval as a function of cell pushing and shoving when forming microcolonies (Angeles-Martinez & Hatzimanikatis, 2021). After the preset simulation time (typically, 10–20 h, or 50,000–100,000 time steps), the final attained individual microcolony biomasses are quantified, and individual microcolony growth rates are fitted from biomass-over-time increases (or back-calculated as occupied colony surface areas). Users can vary the concentration of resources in the model, the production rates of metabolites, and run different interaction scenarios.

To simulate interspecific interactions, we varied production and consumption rates of byproducts (assuming, for simplicity, byproduct ‘groups’; as proposed in (Guex *et al*., 2023)). The influence of mutual or cross-wise utilization of excreted byproducts was then tested by varying the *µ* fraction attributed to the main substrate and to the byproducts, under the restriction that the total *µ* cannot surpass its experimentally observed value (Appendix A). To plot simulations and compare to microscopy images, the simulated cell positions and areas across the surface grid at 20 min intervals were saved as .tif files and segmented using the Python script described above. The cell-agent model was entirely coded in Matlab (version 2021b, MathWorks Inc).

### Metabolite analysis in liquid culture

To characterize appearing metabolites during growth of either species in mono- or coculture, we grew Pve and Ppu in liquid MM with 10 mM succinate in Erlenmeyer flasks (500 ml with 100 ml culture, incubated at 30 °C and 120 rpm rotary movement), and then swapped the cells at mid-exponential phase to continue growing in either their own culture medium or that of the other species. Individual monocultures were grown in biological triplicates to mid-exponential phase (3-6 h after inoculation, measured by culture turbidity – OD_600_ - using a Ultrospec 500 pro spectrophotometer from Amersham Biosciences), after which 90 ml were removed and centrifuged to recover the cells. The supernatant was decanted without disturbing the cell pellet and then further purified from cells by passing over a 0.2-µm membrane filter (ClearLine, 037044). The cell pellet was carefully resuspended in 2 ml sterile saline solution (0.9% NaCl) and its turbidity was measured. The cell-free supernatant was divided in equal volumes in two new sterile Erlenmeyer flasks, one of which was inoculated with its ‘own’ resuspended cell pellet; the other with the resuspended cell pellet from the other species; both targeting a starting OD of 0.005. The cultures were then again incubated as before, and their growth was followed by spectrophotometry.

Aliquots (1.5 ml) for liquid chromatography mass spectrometry (LC-MS) analysis were taken at the start (i.e., uninoculated medium); the mid-exponential phase (i.e., the filtered supernatants before swapping), and at the end of the incubations (i.e., swapped cultures). All samples were purified by filtering across a 0.2–µm membrane filter, then snap-frozen in liquid nitrogen, and stored at –80 °C until analyzed by targeted LC-MS (Agilent 6495 LC-MS QqQ system, conducted at the University of Lausanne, Metabolomics Unit).

### Code availability

The cell-agent surface growth model with description (code in MATLAB vs 2021b) can be downloaded from GitHub (https://github.com/IsalineLucille22/Surface_Chapter_PhD). The Python microcolony segmentation script is available from GitHub (https://github.com/Flamanjaune/PHD-Thesis/tree/main/Part_III/Chapitre_7). All further image analysis code (in R or in MATLAB) with the raw segmented colony areas is available from a single download at Zenodo (to be completed).

## RESULTS

### Extracting growth rates from microcolony area expansion rates

To extract kinetic parameters, we followed real-time formation of microcolonies from individual cells of mono- or cocultures of Ppu and Pve, sparsely seeded on the surface of an agarose patch (1 mm thick and 1 cm diameter). The patch is embedded in a closed microscopy chamber under inclusion of air (Fig. 1A) (Reinhard & van der Meer, 2014). The agarose contains a limiting amount of substrates to control the extent of colony growth (while acknowledging that the agarose itself may also provide some carbon and nutrients for growth – see below). The agarose patches provide the cells with a solid surface to form microcolonies while preserving the diffusion rates of substrate molecules and metabolites close to that in liquid. Dividing cells will expand primarily in the two-dimensional plane, because they are enclosed between the agarose surface and the glass coverslip, facilitating imaging (Fig. 1A). By reducing the amount of primary carbon substrate or increasing the density of founder cells on the patch, the number of divisions per cell is limited and the colonization of the surface remains restricted to single-layered individual microcolonies, which occasionally merge when founder cells fall relatively close to each other. To avoid boundary effects on microcolony growth under influence of a radial oxygen gradient (Supplementary figure 1), we imaged microcolonies in a 3 by 3 mm^2^ area close to the patch centre for consistency.

Since cells are tracked over time by imaging from the onset of the experiment (Fig. 1B), their apparent maximum specific growth rate can be calculated from the segmented colony area expansion during the exponential growth phase for each microcolony (Fig. 1C). To support the empirical observations of colony growth, we developed an accompanying model based on substrate diffusion, uptake and utilization by spherocylindrical cell-agents with similar geometry as either Ppu or Pve cells (Fig. 1D). The model takes into account cell elongation, division and cell-cell pushing during colony development, and recapitulated observed microcolony growth patterns (Fig. 1E). We then further used the model simulations to understand diffusive substrate-growth effects and the nature of emerging interactions in the observed mono- and coculture experiments (see below).

### Microcolony growth rates as a function of added substrate concentration

In order to benchmark the system of surface colonization for Monod kinetics, we first examined the dependency of monoculture growth rates on increasing substrate concentrations. For this we added succinate at increasing concentrations (0.01, 0.05, 0.5, 1 and 5 mM) and seeded cells of either Ppu or Pve individually, with a founder cell density per imaged area (ca. 1.7×10^4^ µm^2^) ranging from 30 to 70 cells for Ppu and from 30 to 150 cells for Pve. Average colony expansion rates increased from 0 to 5 mM succinate, indicative for Monod-substrate dependency (Fig. 2A). Both fitted average µ_max_ and K_S_ on succinate were higher for Ppu (0.63 h^−1^ and 0.023 mM, respectively), than for Pve (0.43 h^−1^ and 0.005 mM). Colony expansion rates did not decrease to zero in absence of any added succinate, indicating that both species can extract some carbon from the agarose medium (Fig. 2A, Supplementary figure 2). By interpolating the fitted Monod curve, we estimated ca. 50 µM of succinate-equivalent carbon source to be available from the agarose. These results thus indicated that Ppu is the faster grower, but Pve potentially has a lower affinity for the substrate under the patch conditions.

**Figure 2.**
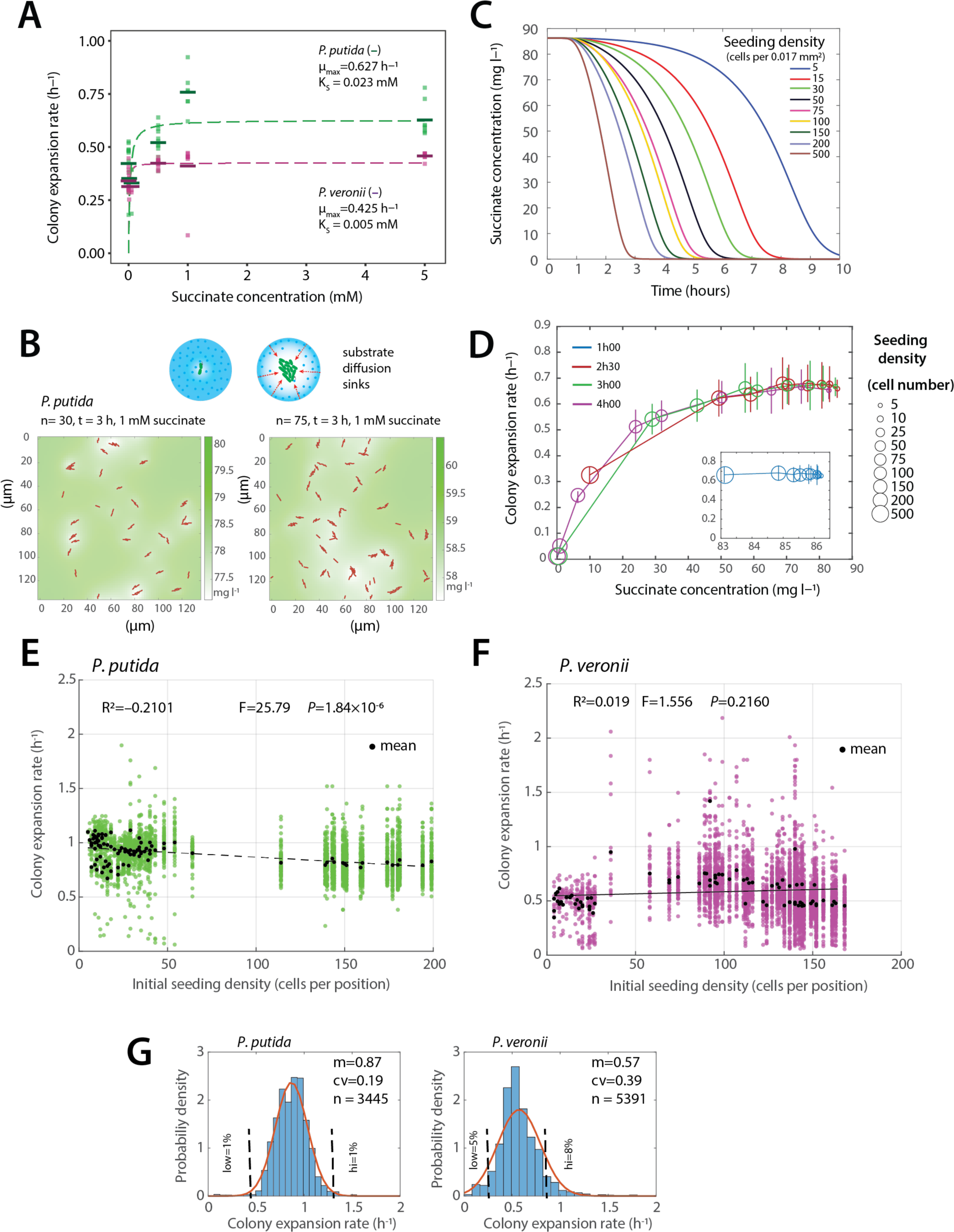
Growth kinetic properties of surface-grown *P. putida* or *P. veronii* founder cells. A) Mean colony expansion rates of *P. putida* or *P. veronii* as a function of succinate concentration in the agarose disks. Horizontal bars, means of average rates across image areas (each consisting of between 15-150 microcolonies, indicated by squares). Dotted lines and values show non-linear Monod fitting to the means. B) Simulated substrate depletion as a function of incubation time and microcolony size in a virtual image area (134 x 134 µm^2^). Cells are shown in red (simulated sizes and orientations). Shades of green correspond to the simulated substrate concentration around the cells, according to the respective color scale on the right. C) Average simulated substrate depletion times (starting from 1 mM succinate) as a function of cell seeding density. D) Mean simulated colony expansion rates with the indicated starting cell densities (circle sizes, to the scale on the right), as a function of substrate concentration, at the indicated incubation time points (colors). Note that at high starting cell density (e.g., 500), the substrate concentration in the patches will have depleted significantly, resulting in a lower measured colony expansion rate (e.g., 0.3 after 2h30). E) Effect of starting cell density (cells per image area position, ca. 0.017 mm^2^) on the measured colony expansion rates of *P. putida* on 1 mM succinate. Black dots are the mean expansion rates within an image area, with green dots being the individual microcolony values. Statistics from linear regression on the means, testing the significance of difference to a zero slope. F) As (E), but for *P. veronii*. G) Distribution of colony expansion rates at 1 mM succinate for *P. putida* and *P. veronii* in monoculture conditions (six independent experiments). n, number of measured microcolonies; m, mean; cv, coefficient of variation (mean µ/standard deviation). Vertical dashed lines indicate the mean ± 0.5×mean, to represent the proportion of colonies with outlier behaviour (low and hi; in percent).

To better understand the substrate utilization by cells on the surface, we simulated Ppu growth at different cell seeding densities using the observed colony expansion rates at 1 mM succinate, and calculated the average remaining substrate concentrations in the agarose patch as a function of incubation time (Fig. 2B). This indicated, as expected, that the denser the amount of seeded cells on the surface, the faster the substrate is depleted (Fig. 2C). In other words, this would mean that at very high starting cell densities (>200 per image area), the time window to accurately measure the exponential growth rate after seeding the cells would be less than 2 h (Fig. 2D). In our further protocol, therefore, we targeted less than 150 seeded cells per unit of imaged area, and started the imaging maximally 30 min after seeding the cells. Based on these results, we used 1 mM succinate in the following experiments, which we concluded is not maximum growth rate-limiting and would permit substrate competition, while still restricting multi-layered colony growth.

In accordance with model predictions, the measured mean individual colony expansion rates at 1 mM succinate in six independent monoculture experiments remained relatively consistent across a range of starting cell densities (5 – 200 per imaging area of 0.017 mm^2^). Ppu colony expansion rates decreased slightly but statistically significantly as function of increasing founder cell density (Fig. 2E; R^2^= –0.2100, p-value = 1.84×10^−6^), whereas those of Pve did not (Fig. 2F; R^2^= 0.0193, p-value = 0.2160). The observed colony expansion rates were less variable for Ppu (mean = 0.87 h^−1^, coefficient of variation = 0.19, Fig. 2G) than for Pve (0.57 h^−1^ and a cv of 0.39), with 1% low and high Ppu microcolony outliers versus 5 and 8% for Pve (Fig. 2G, low and high defined as below or above the mean expansion rate ± 0.5 × the mean). As the substrate is homogenously present in the patches, this suggests an inherent phenotypic variability among founder cells, determining their reproductive success.

### Growth kinetic heterogeneity among individual microcolonies

Despite different expansion rates, all colonies in the imaged areas entered stationary phase at almost the same moment (visible from the arrest of colony area increase), whereby Ppu in mono-culture (Fig. 3A) reached stationary phase sooner than Pve (Fig. 3B; note that a further slow increase is detected at t>300 min, which is due to continued expression of the fluorescent marker in non-dividing Pve-cells that inflates the segmented colony area). Growth arrest of individual microcolonies is a consequence of one or more factors becoming growth-limiting. Their near-simultaneous stalling, despite being surrounded by appreciable empty space (Fig. 1E), suggests almost homogenous depletion of growth factors at the scale of the imaged areas. Therefore, even though larger colonies tend to locally deplete substrate concentrations (as indicated by simulations in Fig. 2B), molecular diffusion would rapidly counterbalance and equalize substrate concentrations at the imaged area scales (Fig. 2C and D). As a consequence of the near simultaneous growth cessation, the microcolonies reached different stationary phase sizes. The mean microcolony sizes for both Ppu and Pve decreased from 1 to 0.5 and 0 mM succinate (with same founder cell densities), as expected (Fig. 3C). Simulations showed, however, that individual microcolony sizes at stationary phase are largely dependent on the founder cell density (Fig. 3D, E), and their absolute sizes are, therefore, only of limited value to judge differences in biomass productivities as a result of interspecific interactions (such as substrate competition; see below).

**Figure 3.**
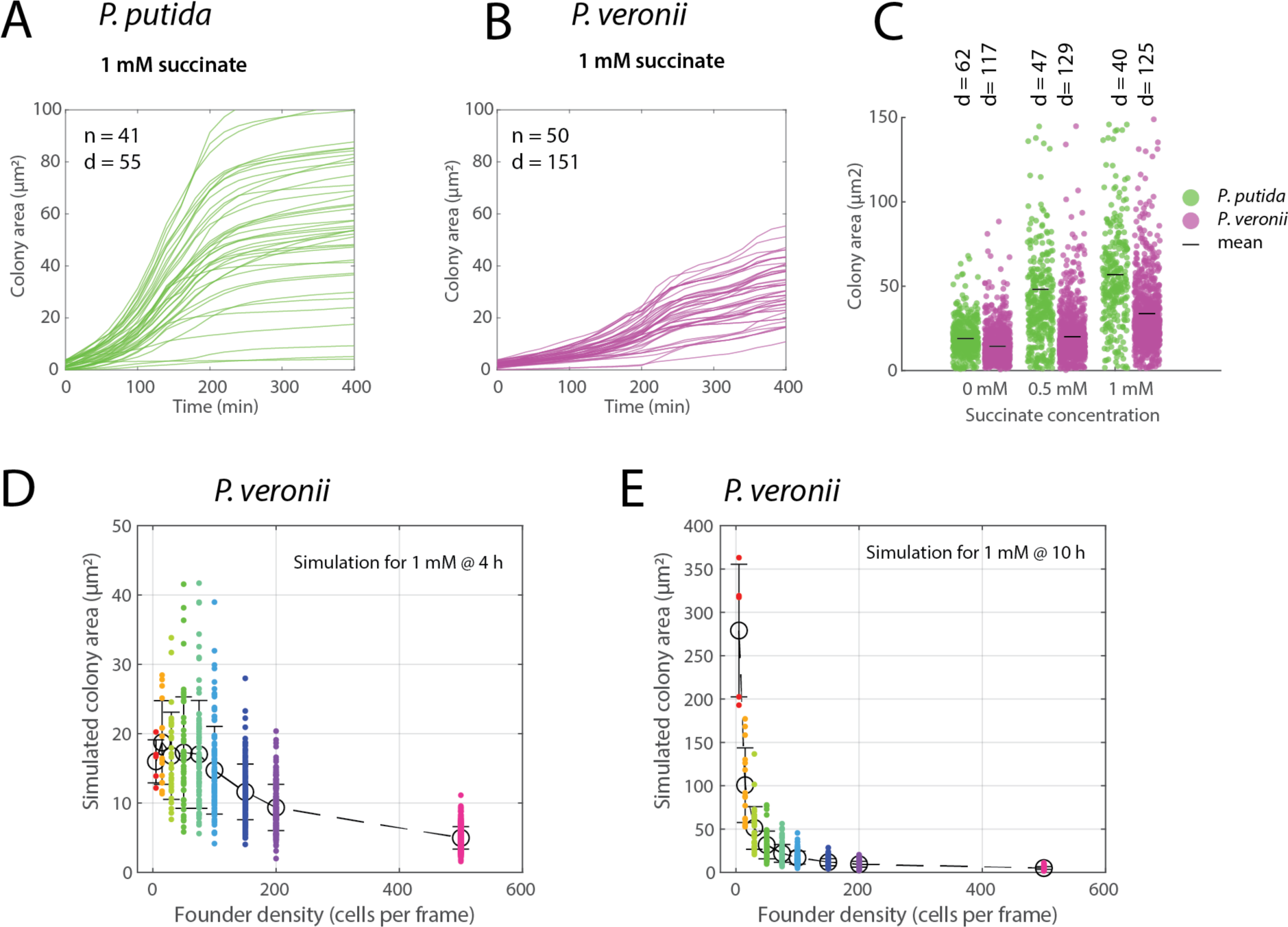
Microcolony productivity variation as a function of substrate concentration and founder cell density. A) and B) Microcolony growth and steady state area variation of *P. putida* (A) or *P. veronii* (B) with 1 mM succinate on a single imaged area as function of time, measured from area occupied by the fluorescent cells (as in Fig. 1B). Note how all colonies arrest growth at approximately the same time. C) Observed microcolony size variability in stationary phase as a function of succinate concentration. Each dot is an individual microcolony. d, mean founder cell density (cells per frame). D) and E) Simulated microcolony areas of *P. veronii* on 1 mM succinate after 4 h (D) and 10 h (E) as a function of founder cell density. Circles represent the mean of individual simulated microcolonies (colored dots), with bars representing the standard deviation.

To further understand the cause of colony size variability, we examined three possible factors: the number of neighbors at start, the apparent colony expansion rate, the lag phase of the founder cells (here taken as the time until first doubling of the initial cell area), and the area of Voronoi tesselation per founder cell (Chacon *et al*., 2018). The number of nearby neighbors (scored as the number of neighboring founder cells in circles with increasing radii) correlated negatively with the final observed microcolony size, both for Ppu (Fig. 4A) and for Pve (Fig. 4B). Significant but weak correlations were found between the tesselated Voronoi area at start and the final attained microcolony areas for both Ppu and Pve (R^2^= 0.3069 and 0.4429, resp.; Supplementary figure 3). However, there was no correlation of the number of neighbors and the observed colony expansion rates (Supplementary figure 4). This means that at the start, all founder cells perceive sufficient substrate influx to grow at maximum exponential rates. Most founder cells taken from exponentially growing precultures did not exhibit any apparent lag phase once they were deposited on the surface and the imaging had started, because their measured colony expansion rates were inversely proportional to the time of their first doubling (Fig. 4C and D; *expo*, exponential phase precultures). In contrast, Ppu founder cells prepared from stationary phase liquid suspended cultures showed ca. 10% of cells with a detectable longer lag phase than expected from their colony expansion rate (Fig. 4C), whereas this was less than 1% for Pve (Fig. 4D). Interestingly, however, Ppu cells with longer lag times still displayed fast maximum growth rates. Collectively, these results indicated that the variable starting maximum growth rates of individual cells are predetermined at the time of seeding, perhaps as a consequence of preculturing history and cell phenotypic variability. Starting (maximum) growth rates of individual cells are not necessarily decreased by longer lag times.

**Figure 4.**
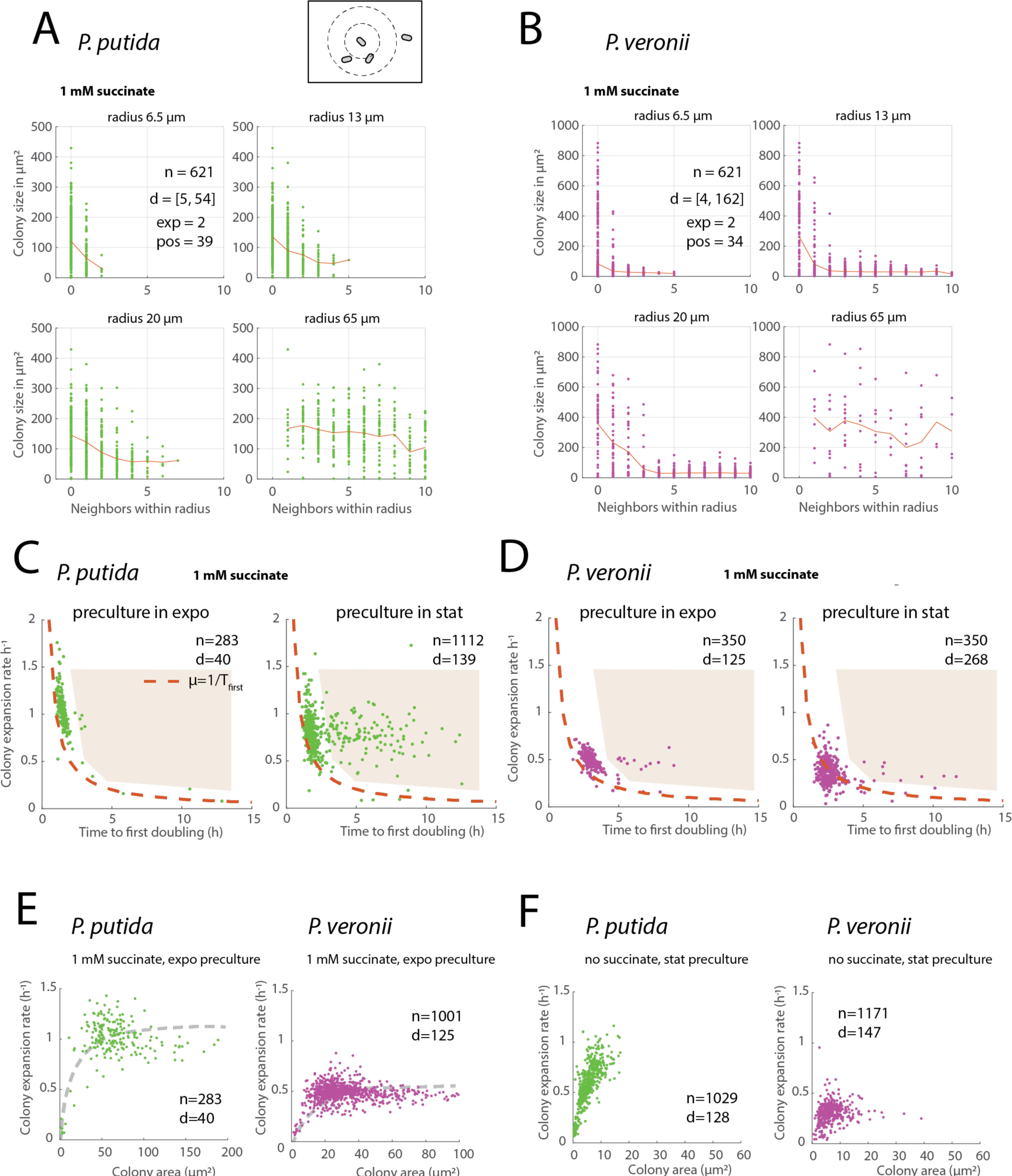
Dependencies of microcolony growth properties on available surface area. A) and B) Relationship between attained colony areas (individual dots) and area occupation by neighbouring kin, calculated as the number of colonies within a circle of diameter *x* around each cell (as depicted in the cartoon). Colony data from patches with 1 mM succinate in monoculture. n, number of plotted microcolonies; d, range of founder cell densities; exp, number of independent experiments and pos, imaged positions. Red lines connect the means. C) and D) Relationship between measured colony expansion rate and the time to first doubling (as the sum of the fitted lag time and the inverse of the measured colony expansion rate) for *P. putida* (C) or *P. veronii* (D) monocultures on 1 mM succinate, seeded either from exponentially growing precultures (expo), or stationary phase cultures (stat). n, number of microcolonies in the plotted datasets (*P. veronii* subsampled to 350); d, mean founder cell density per image area. Dotted line shows the expected time to first doubling without lag phase (as 1/colony expansion rate). Shaded areas indicate cells with extended lag times. E) and F) Stationary phase colony areas as a function of measured colony expansion rates for *P. putida* or *P. veronii* at 1 mM succinate, with precultures from exponential phase (E) on the same substrate, or from stationary phase cultures seeded on patches without succinate added (F). Dotted lines show indicative dependency.

As a consequence, stationary phase colony areas were not well explained by measured expansion rates (Fig. 4E). That this is not a general rule is shown by cells taken from stationary phase precultures on succinate and seeded on surface without succinate (therefore only being able to profit from residual carbon in the agarose, Fig. 4F). In such case, the final microcolony sizes do correlate to the individual colony expansion rates (Fig. 4F). In summary, monoculture (maximum) growth rates of individual founder cells and their variability seem mostly predetermined by preculture conditions and stochastic variation, whereas individual colony productivity (i.e., their stationary phase size) is to some extent dependent on the number of neighbors and the available area, as expected (Chacon *et al*., 2018). Additionally, this suggests that faster or bigger growing microcolonies lead to a substrate ‘sink’, which locally biases diffusion towards them.

### Substrate competition in cocultures leads to growth rate reduction and local colony size effects

Having set the boundary conditions on microcolony growth rates and yields from monoculture data, we next compared colony expansion differences to cocultures of Ppu and Pve. We first focused on conditions where we expected both species to engage in competition for the same primary substrate (succinate). Since Ppu grows faster than Pve on succinate (Fig. 2E), our expectation here was that Ppu would outcompete Pve growth as seen in liquid cultures (Guex *et al*., 2023). Unexpectedly, microcolony expansion rates of both Ppu and Pve were lower in coculture than in monoculture, but coculturing had no or little effect on measured lag times (Fig. 5A). The coculture-dependent decrease of colony expansion rates was consistent across multiple different experiments, even at different succinate concentrations and seeding densities, with an average decrease of 16.4% for Ppu and 14.9% for Pve (Fig. 5B, n= 10 and 8, and p-values of 6.39×10^−5^ and 0.0026, respectively). Individual microcolonies of either strain were on average smaller in co-than in corresponding monocultures (Supplementary figure 5). In terms of global productivity (e.g., summed stationary phase colony size for each of the strains per imaged area position), Pve lost 90% in coculture with Ppu compared to monoculture, whereas Ppu lost on average 27% (Fig. 5C). There was no significant difference in summed productivity of both species in coculture and that of Ppu in monoculture (p=0.1776; Fig. 5C). The competitive loss factor on growth rate for Ppu (as the relative difference compared to monoculture growth) thus equalled 1–0.164 = 0.836 and on productivity 1–0.27=0.73, whereas that of Pve equalled 1–0.149 = 0.871 on growth rate and 1–0.90 = 0.10 on productivity. Cell counts measured by flow cytometry across the whole patch at stationary phase showed similar competitive loss for Pve in the coculture compared to its monocultures (0.10; Fig. 5D), but no significant difference for Ppu productivity in the mixture compared to its monocultures (p=0.3075, n = 10).

**Figure 5.**
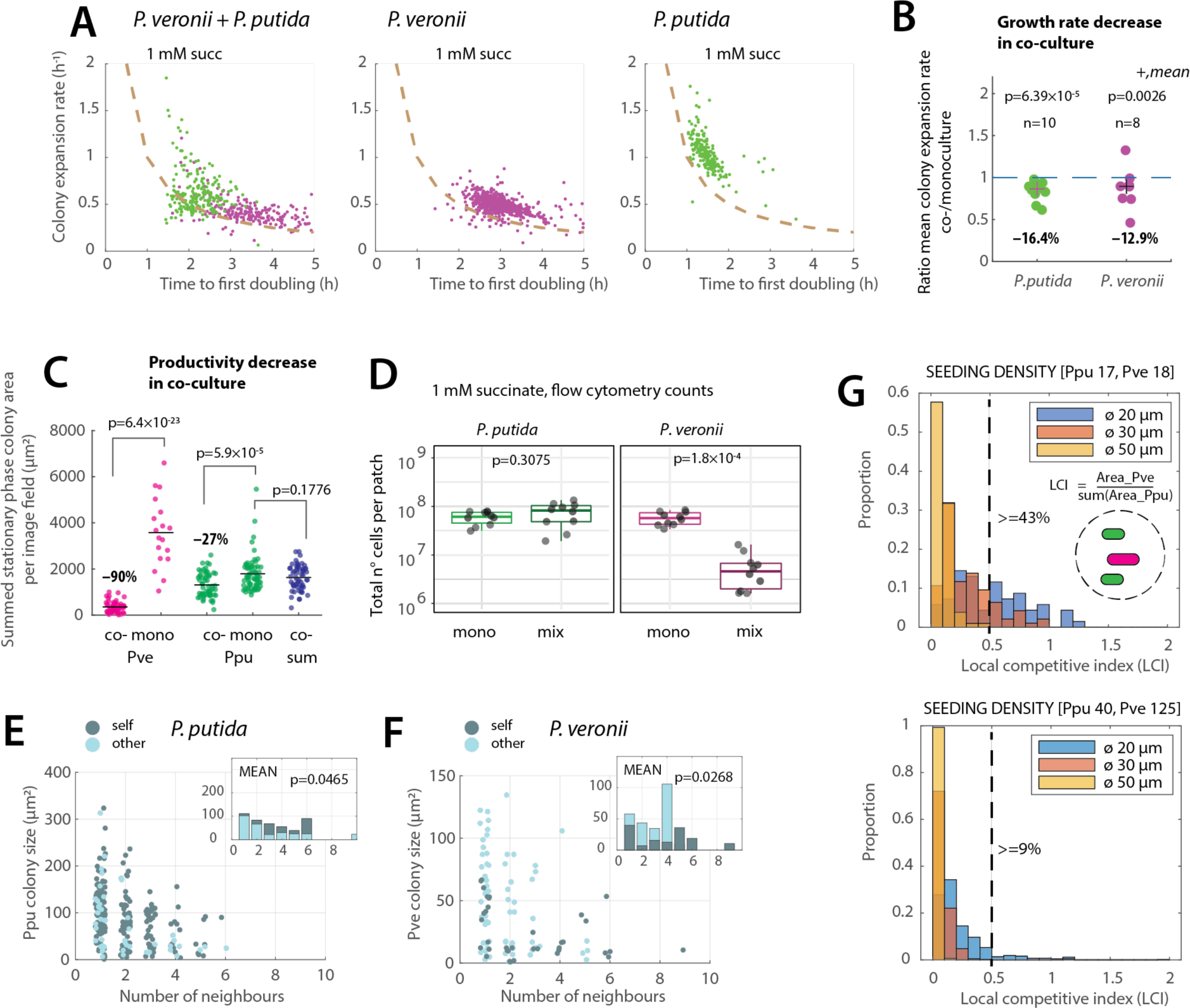
Effect of coculturing on microcolony growth properties. A) Colony expansion rates for *P. putida* (green dots) or *P. veronii* (magenta) in monoculture, or in coculture on 1 mM succinate, as a function of time to first doubling (h; being the inverse of the measured colony expansion rate plus fitted lag time). Dotted lines show the expected relation in absence of lag time. Precultures were taken from exponential phase on the same substrate. B) Difference of mean microcolony expansion rates in coculture compared to monoculture (n, number of experiments; p-values from unpaired t-testing). C) Productivity decrease in coculture compared to monoculture, taken as the sum of microcolony stationary phase areas per imaged field. Dots represent technical replicates (image fields) across experiments with the same succinate concentration. P-values from unpaired t-testing. Co-sum, per image field summed areas of both *P. putida* and *P. veronii.* D) Total number of cells on patches of mono- or cocultures, grown with 1 mM succinate, determined by flow cytometry of washed suspensions. E) and F) Effect of neighbouring *self* or non-kin (*other*) microcolony density within a 30–µm circle radius around the cell on its final attained microcolony size (Ppu, *P. putida;* Pve, *P. veronii*). Inset (bars) shows the means at the individual neighbour values. Note how *P. putida* microcolonies tend to be smaller with surrounding *P. veronii* colonies, and *P. veronii* colonies tend to be bigger when surrounded by *P. putida*. P-value from Wilcoxon single-tail rank-sum test. G) Proportion of *P. veronii* microcolony areas compared to that of the sum of *P. putida* areas within circles of the indicated diameter (i.e., local competition index or LCI). The dotted line indicates the global mean proportion of *P. veronii* biomass compared to *P. putida* (i.e., as from panel C) + 2 × the standard deviation, and the number refers to the percentage of *P. veronii* microcolonies with LCI above the threshold for the case of a diameter of 20 µm.

To estimate competition at a local scale, we compared microcolony areas of either species surrounded by only colonies of the same kin (Fig. 5E&F, *self*) or only of the other species (*other*). Interestingly, this indicated that Ppu colonies surrounded by Pve are smaller than those surrounded by its own kin (Fig. 5E, inset; p=0.0465, Wilcoxon ranksum test), whereas Pve colonies exclusively surrounded by Ppu were slightly bigger than those surrounded by their own kin (Fig. 5F, inset; p=0.0268). This suggests that Pve can locally benefit from Ppu, perhaps through metabolite cross-feeding, as previously suggested (Guex *et al*., 2023). Finally, we expected that spatial and stochastic growth kinetic variations might enable Pve to dominate the competition with Ppu in sparse instances, but only at a local scale (Fig. 5G). Indeed, one can see that, depending on the seeding density, between 9 and 43 % of Pve microcolonies grow bigger than expected from the global competitive effects (= 0.1), but these effects are lost at a larger scale (ranges of 30 and 50 µm, Fig. 5G).

To place the effect of primary substrate competition into perspective, we repeated the same experiment, but with an exclusive substrate for each of the species such that they would become indifferent for each other. The best combination to achieve this and which abolished competition in liquid culture (albeit not perfect) used putrescine as selective substrate for Ppu and D-mannitol for Pve (Guex *et al*., 2023). Despite the intended indifference, some 10 % of Pve colonies showed diauxic growth, indicating they may be able to use putrescine (at a later stage, Fig. 6A). As intended, the productivity of both species on the combination of putrescine and D-mannitol in coculture was less drastically affected (Fig. 6B, loss factors 0.61 and 0.67 for Ppu and Pve, resp.) than with succinate (0.73 and 0.10; Fig. 5E), although the summed stationary phase productivity in the coculture was less than the sum of the monoculture productivities (Fig. 6B). This indicates that also under substrate indifference conditions both species do not grow completely independently from each other. Individual maximum growth rates observed for Ppu (taken at the first 4 hours) were on average two-fold increased in presence of Pve (Fig. 6C and D), but no different for Pve in presence or absence of Ppu (Fig. 6D). In presence of Ppu, however, the variability among growth rates of Pve microcolonies increased (Fig. 6C). In contrast to what was observed for succinate, the final attained sizes of Ppu microcolonies on patches with D-mannitol and putrescine, surrounded exclusively by Pve (within a radius of 10 µm), were not different than those surrounded exclusively by other Ppu microcolonies, and vice versa (Fig. 6E). This suggested that, locally speaking, there was no evidence for competitive interactions under the substrate indifference scenario.

**Figure 6.**
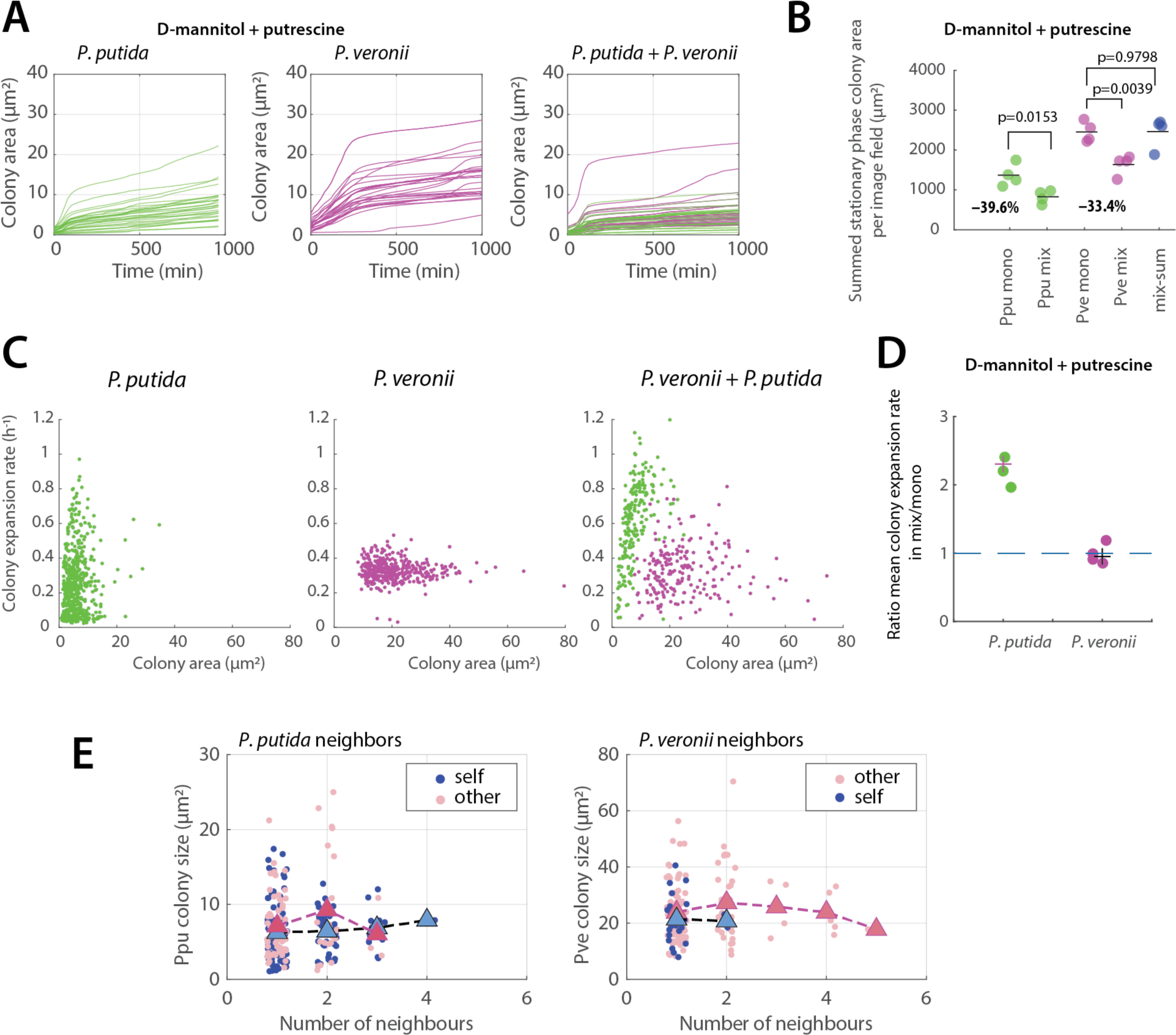
Coculture behaviour under substrate indifference conditions. A) Microcolony growth of *P. putida* or *P. veronii* separately or in mixture on D-mannitol and putrescine. Only the first increase is used to calculate the maximum specific growth rate (i.e., typically between 0 and 200 min). B) Stationary phase productivity difference in mono- or coculture growth (productivity is the sum of microcolony areas per image field, here as individual dots). C) Colony expansion rates as a function of stationary phase colony area. D) Ratio of mean growth rates in co-versus monoculture for either *P. putida* (PPU) or *P. veronii* (PVE). E) Effect of neighbouring *self* or non-kin (*other*) microcolony density within a 30– µm circle around the cell on the final attained microcolony size. Triangles show the means of the colony size distributions.

### Surface coculture growth simulations underscore metabolite-driven interactions

To better understand the potential causes of observed coculture effects on kinetics of microcolony growth, we simulated different interspecific interaction scenarios. As for the monocultures, coculture simulations were started with randomly placed founder cells of either species or else with starting positions matching those of the experiments. Cells of either species were given growth kinetic properties based on their monoculture experimental values, plus a random variation equal to the observed standard deviation. We did not exhaustively investigate parameter space effects, but concentrated on simulating a number of potential credible scenarios (Fig. 7A). Direct competition for the single added substrate (succinate), was not able to explain the observed reduction in growth rates; neither was a simulation scenario with sharing of the same metabolites (Fig. 7B). In contrast, simulating both cross-feeding (i.e., production and uptake of different metabolites) or cross-feeding with interspecific inhibition resulted in a reduction in growth rates of both species (Fig. 7B). As expected, all scenarios predicted a lower productivity of Pve compared to Ppu, and all but direct substrate competition caused a reduction in the (relative) productivity sum (Fig. 7C). Inclusion of a mutual inhibition resulted in the largest reduction in the productivity sum (Fig. 7C). The variation of simulated individual colony sizes and their maximum expansion rates for the cross-feeding and inhibition scenarios captured the main trends observed in the experimental data (Fig. 7D). However, removing from the simulations either the variation in maximum growth kinetics or variation in lag phase of starting cells, yielded much less variation in colony expansion rates, which is not in agreement with experimental observations (Fig. 7D). This underscores, therefore, the crucial effect of inherent differences in growth kinetic properties of founder cells on the productivity of their microcolony descendants.

**Figure 7.**
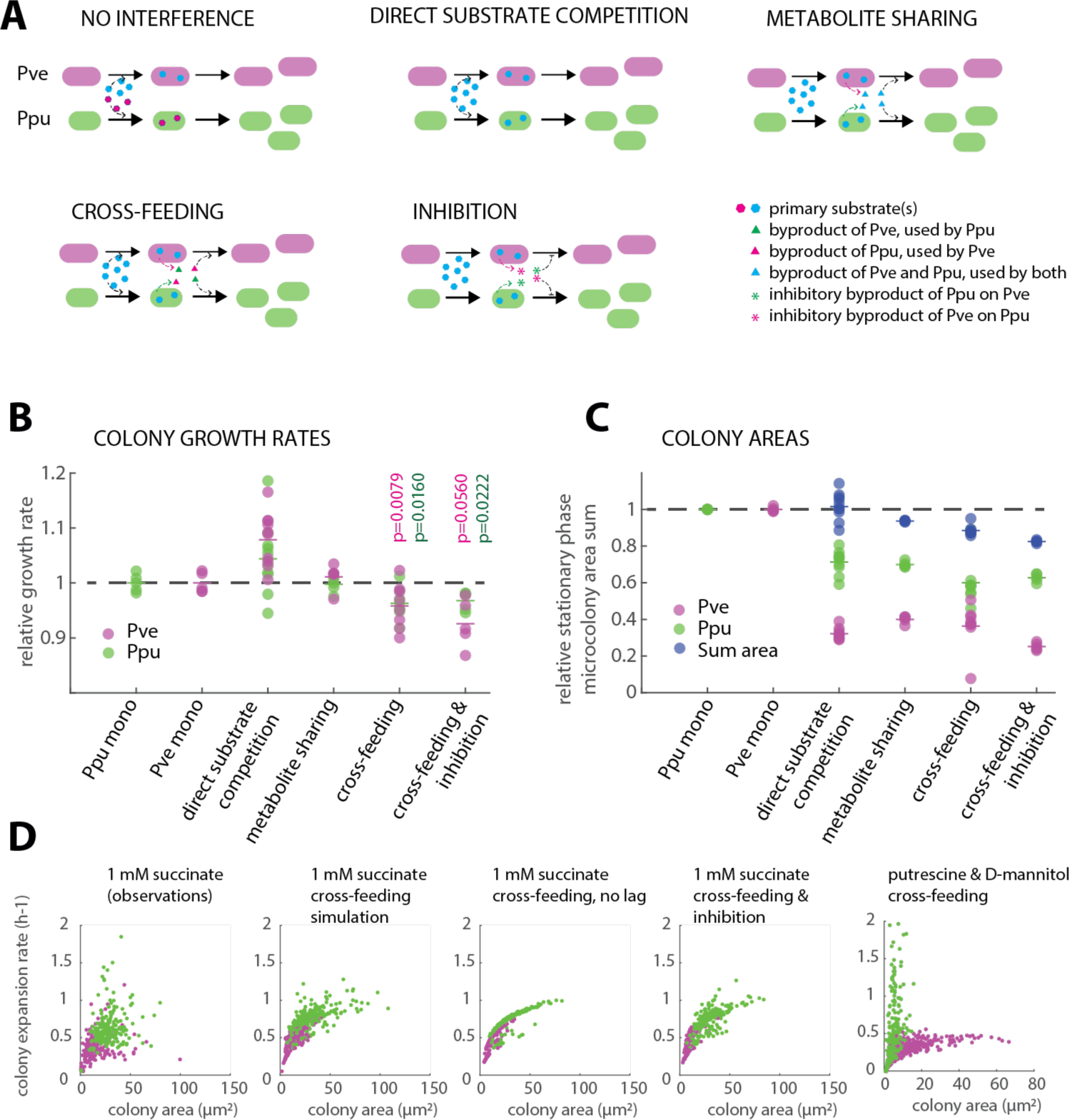
Simulated interaction scenarios explain reduced growth rates in cocultures. A) Different tested scenarios of cross-feeding or inhibition between *P. putida* (Ppu) and *P. veronii* (Pve). B) Relative growth rates of either *P. putida* or *P. veronii* in competition scenarios compared to their monoculture simulations (n=5–7 simulations). P-values correspond to unpaired t-tests between co- and monoculture simulation replicates. C) as B, but for the final attained microcolony area sum (per simulated image field). Sum area, sum of *P. veronii* and *P. putida* colony areas in that simulated image field. D) Relation between colony expansion rate (simulated or experimental) and final attained microcolony area.

### Extensive production and sharing of metabolites in competition

To support the suggestion that differential metabolite production and uptake can determine interaction outcomes, we repeated Ppu and Pve mono- and coculture growth in liquid suspension to facilitate targeted metabolomics analysis. No significant effects on Ppu or Pve growth rates could be detected after swapping their (exponential phase) culture medium to that of the other species (Fig. 8A, Supplementary figure 6), although Pve with Ppu-supernatant showed a shorter delay in resuming growth (Supplementary figure 6). The excreted metabolites in pure cultures were very similar among both species, and all accumulated over time, except dihydrouracil (Fig. 8B). Ppu notably produced higher concentrations of (5’-)deoxyadenosine, cis-aconitate and homocysteine than Pve, whereas Pve produced more alpha-ketoglutarate, cyclic-GMP, ethanolamine, N-acetylputrescine and salicylate than Ppu (Fig. 8B). The sum of the peak areas of the produced metabolites accounted for 25.4% of the initial succinate concentration (in peak area) for Ppu, and 23.5% for Pve. Despite a number of obvious differences in mono-culture metabolite production, both Ppu and Pve showed very few significant differences in the utilization and production of metabolites starting from their own exponential phase supernatant or from that of the other species (Fig. 8C). At a cutoff of twofold difference and adjusted p-value of 0.05, only Ppu produced more cyclic-GMP, deoxyguanosine, N-acetyl leucine, and thymidine with Pve’s supernatant than with its own (Fig. 8C). These results thus indicated that both Ppu and Pve produce very similar metabolites, but can reciprocally reutilize those which are specifically produced by the other species.

**Figure 8.**
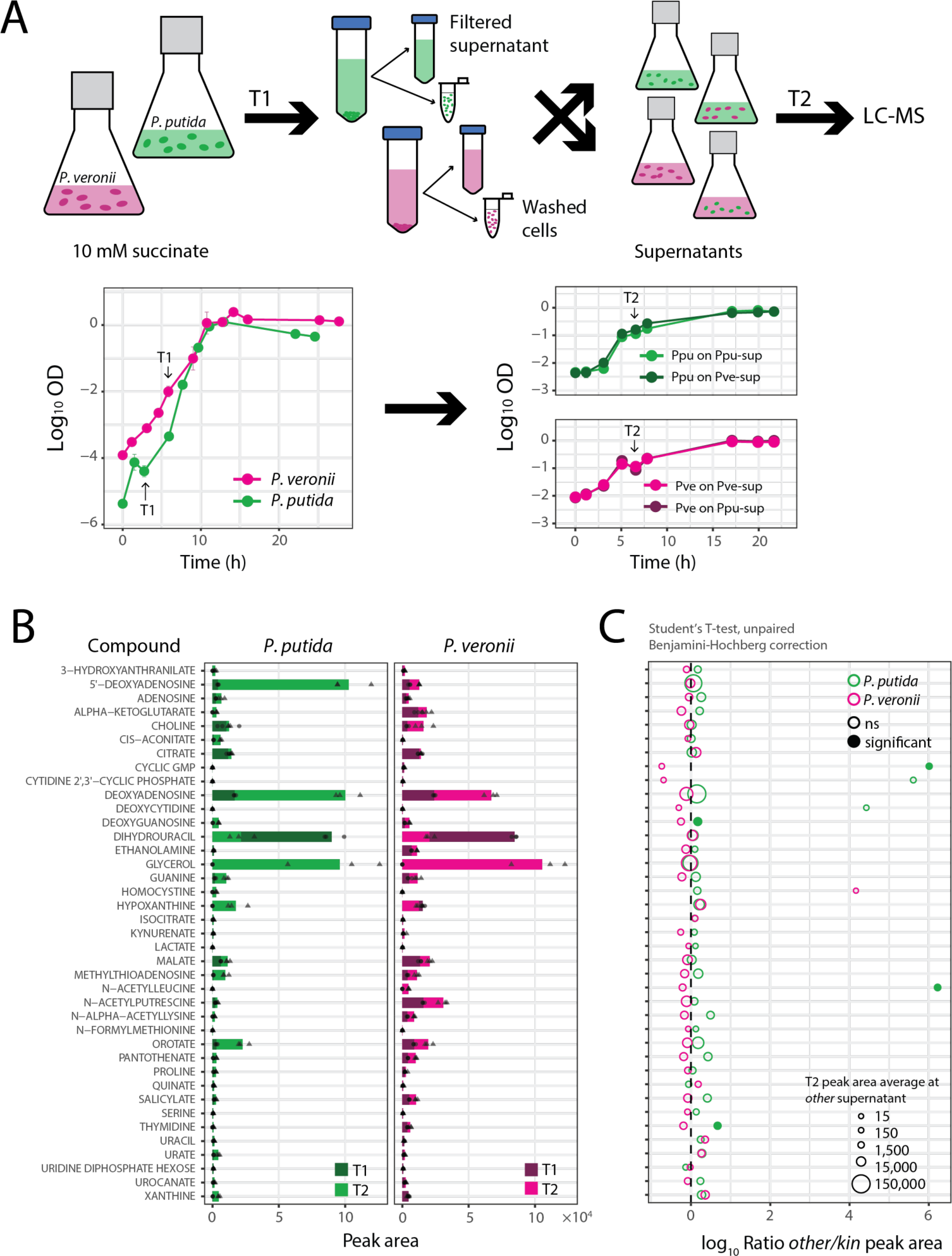
Exometabolite analysis of growing mono- and cocultures. A) Growth of *P. putida* and *P. veronii* in monocultures on 10 mM succinate and on reciprocal cell-free supernatants harvested from exponential cultures of either strain (for more detailed time series and growth rate determinations, see Supplementary figure 6). Dots represent the means of independent biological triplicates. T1, T2: sampling and swapping times. B) Targeted detected metabolites in monoculture supernatants of either strain in exponential (T1) and early stationary phase (T2). Bars show the mean detected peak area from biological triplicates, with circles (for T1) and triangles (for T2) showing individual values. C) Log_10_ ratio of the compounds’ T2 areas in incubations of the *other* species’ and its own supernatant (*kin*). Significance of difference by paired t-test corrected for multiple test correction (as filled circles).

## DISCUSSION

We successfully showed that a simple microcolony growth platform can be expanded from mono-(Eijlander & Kuipers, 2013, Koutsoumanis & Lianou, 2013, Nghe *et al*., 2013, Jung & Lee, 2016, Sankaran *et al*., 2019) to cocultures to parametrize growth kinetic and interspecific interaction effects. Similar setups to study paired interactions are so far extremely rare (Niggli *et al*., 2021) (Laffont *et al*., 2024), but, as we demonstrate, have a high potential for producing rich data sets that encompass both individual and local variability as well as global growth interaction effects. In that sense, the microcolony platform is versatile, quantitative and extrapolates across scales. The platform is easily adaptable to different growth conditions or spatial heterogeneities, and can potentially be scaled to higher order mixed cultures by colony phenotypic recognition (Marcoux *et al*., 2014, Paquin *et al*., 2022, Doh *et al*., 2023) instead of genetically encoded fluorescence to differentiate the strains.

How well does the microcolony platform capture and scale the imposed interaction effects (substrate competition and indifference)? The loss of individual biomass formation for Ppu or Pve under substrate competition in co-compared to monocultures as observed for the entire patch-population (e.g., Fig. 5D) was similar to that measured previously in liquid suspended cultures (Guex *et al*., 2023). As expected from its higher maximum specific growth rate on succinate, Ppu globally outcompeted Pve both for surface- and liquid grown cocultures. The dominance of Ppu over Pve, and the loss of individual biomass in cocultures was also reflected in the decrease of global mean individual stationary phase microcolony areas (Supplementary figure 5), and the summed colony areas per image field (Fig. 5C). In contrast, the relation breaks down at the level of individual microcolony areas, because their size variability is largely dependent on random positioning, general seeding density effects (Fig. 3D&E and Supplementary figure 3) and the phenotypic variation of founder cells (Fig. 4E&F). At local scales, however, the competitive disadvantage for Pve can be overturned, and the stochastic positioning and founder cell phenotypic variations can result in better than expected Pve microcolony growth. For example, in case when single Pve colonies are surrounded by multiple Ppu nearby (Fig. 5E&F). The frequency of overturned competitive effects is low but non-negligible (Fig. 5G), and may contribute to a better than expected proliferation of poorer competing species.

A more unexpected finding with the microcolony coculturing platform was the consistent measured decrease in maximum specific growth rates of both Ppu and Pve by ca. 15% in surface-grown cocultures compared to monocultures at the same seeding densities (Fig. 5A&B). This is not *a priori* expected from Monod-theory, which dictates that initial maximum specific growth rates should be solely dependent on prevailing substrate concentrations, which we showed here (Fig. 2) are not limiting for both Ppu and Pve (at least not during the duration for measuring the colony expansion rates). This leads us to conclude that competitive interactions are not only detectable through yield loss, but also from a decrease in maximum specific growth rate. Our control experiment to generate indifference growth conditions indeed alleviated the observed reduction in growth rates (Fig. 6D) and the neighborhood effect (Fig. 6E), but was not completely orthogonal since both strains lost some biomass production in co-versus monocultures (Fig. 6B).

The question remains as to which mechanism explains growth rate reductions under competition. Model simulations with the coculture agent-based model suggested that neither primary substrate competition nor metabolite sharing can explain maximum growth rate reduction (Fig. 7B). The only two likely scenarios that could yield growth rate reduction are the production of some non-kin targeted inhibitor compound, the utilization of a major metabolite that is not used by the producer strain, or a combination of both. Indeed, by analyzing mono- and coculture swapped supernatants of the strains grown in liquid suspension, we found one compound which is exclusively produced by Ppu but not accumulating in Pve cultures (5’-deoxyadenosine; Fig. 5B&C), and a few minor ones for the reciprocal direction (e.g., c-GMP, N-acetylputrescine). The LC-MS targeted analysis confirmed that both strains, as assumed more in general (Goldford *et al*., 2018, Pacheco *et al*., 2019) are rather leaky, and efflux a large variety of metabolites during exponential growth. In contrast, liquid suspended culturing of either strain in reciprocal exponential phase cell-free supernatant could not recapitulate the loss of growth rates, which may therefore specific for the spatial setting. We can thus assume that regular monoculture growth constitutes a combination of direct substrate metabolism leading to new cell biosynthesis, and conversion of substrate to intermediates, some of which leak out but then are taken up again by the cells in culture. In coculture, these intermediates may be partly consumed by the competing species to the detriment of the producer (Guex *et al*., 2023). Mathematically speaking, this can be approached by a summation of multiple growth rates that combine into a single maximum specific growth rate (Supplementary data 1). In a spatial context, the efflux of metabolic intermediates would lead to microcolonies becoming net producers and effectively losing carbon through radial diffusion, from which nearby other competing species may profit. This has been seen before for a binary couple of engineered amino acid auxotrophies in *E. coli* under microfluidic growth (Dal Co *et al*., 2020), but our data suggest this should be a much more general phenomenon that can lead to growth rate reduction and locally overturning of interaction effects (as in Fig. 5E&F).

Most bacteria in natural habitats are dependent on a spatial nutrient context, originating from gradients, substrate flow and diffusion, or substrate dissolution – either chemically or through activities of other microorganisms. In this sense, the agarose patch reflects a dynamic nutrient habitat, where initially all substrate is dissolved homogenously, but gradients rapidly arise as a consequence of cell growth and metabolism. Curiously, we observed that the variation in reproductive success (i.e., the final microcolony sizes) is poorly explained by the variation in measured colony expansion rates, but rather depends on the variation in numbers of neighbouring cells in the available space. This seems intuitive, but, given the fast substrate diffusion rates in the agarose patch, one would think that the same substrate concentration prevails anywhere at any time. However, even microcolonies located in relatively empty areas can remain small and the relation between tesselated Voronoi area at start and final microcolony sizes is relatively weak (Supplementary figure 3). We postulate, therefore, that microcolony expansion is influenced by the biased substrate diffusion as a result of more rapid net substrate uptake by larger microcolonies (Fig. 2B). The net effect of this would be that faster and bigger growing microcolonies tend to intake proportionally more substrate than smaller ones.

In conclusion, the direct quantification of growth kinetic parameters from surface-grown microcolonies in mono- and coculture permits detecting global as well as local interaction effects, which can be used to feed Monod-based models that correctly describe surface coculture growth. The microcolony growth platform is thus a useful expansion to transduce scales in community growth behaviour and link to physiological theory.

## ACKNOWLEDGEMENTS

This work was supported by the Swiss National Science Foundation (Sinergia program, grant CRSII5 189919/1 to J.M. and C.M.), SystemsX.ch grant 2013/158 (Design and Systems Biology of Functional Microbial Landscapes “MicroScapesX” to J.M.), and by the National Centre in Competence Research (NCCR) in Microbiomes (grant number 180575 to J.M.). We thank Simon van Vliet for helpful critical comments on the manuscript.

## AUTHOR CONTRIBUTIONS STATEMENT

T.M.T., M.D., I.G., C.M. and J.M. conceived the studies and designed experiments. T.M.T., M.D. and E.S-L. conducted microcolony growth experiments. I.G. and C.M. developed the surface-growth cell-agent based model. I.G. scripted the model code, and I.G. and J.M. performed model simulations. X.R. and H.T. conceived the microcolony image analysis and codes. T.M.T., M.D. and J.M. analyzed microcolony data. T.M.T., I.G. and J.M. analysed output data, and wrote the draft manuscript. All authors gave input, verified and corrected the written manuscript. C.M. and J.M. acquired funding and coordinated the work.

## COMPETING INTERESTS STATEMENT

The authors declare no competing interests.

## Notes

https://github.com/IsalineLucille22/Surface_Chapter_PhD

https://github.com/Flamanjaune/PHD-Thesis/tree/main/Part_III/Chapitre_7

### Competing Interest Statement

The authors have declared no competing interest.

