## Supplementary information for "Inferring Bacterial Interspecific Interactions from Microcolony Growth Expansion"

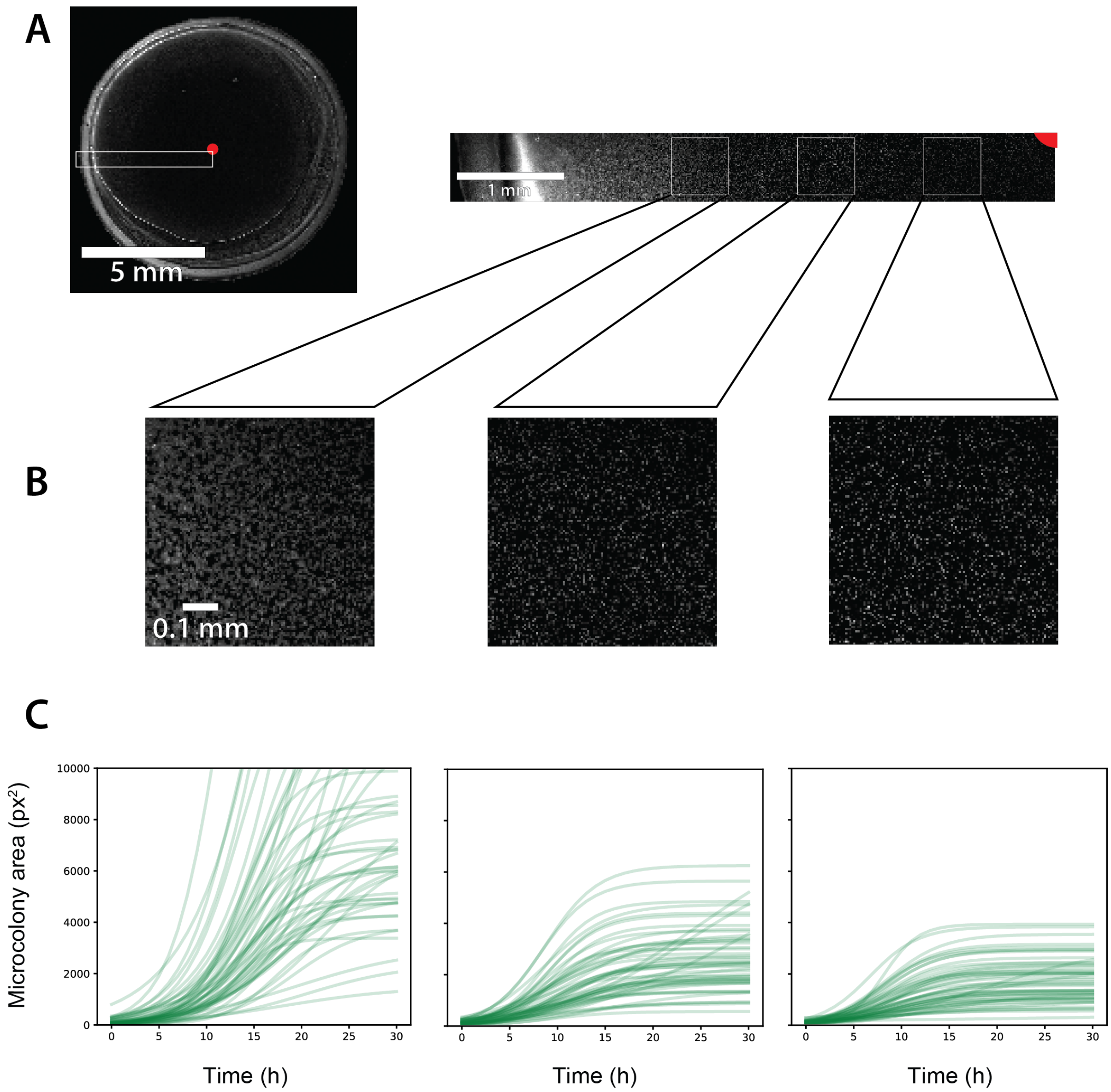


**Supplementary figure 1. Microcolony size distribution as a function of radial distance over the agarose patch.**

(A) Low magnification image of *P. putida* fluorescence distribution over the whole patch and within a radial transect. (B) Higher magnification of and (C) microcolony growth in the squares in the transect in (A), showing larger microcolony areas near the patch edges, suggesting radial oxygen diffusion effects.

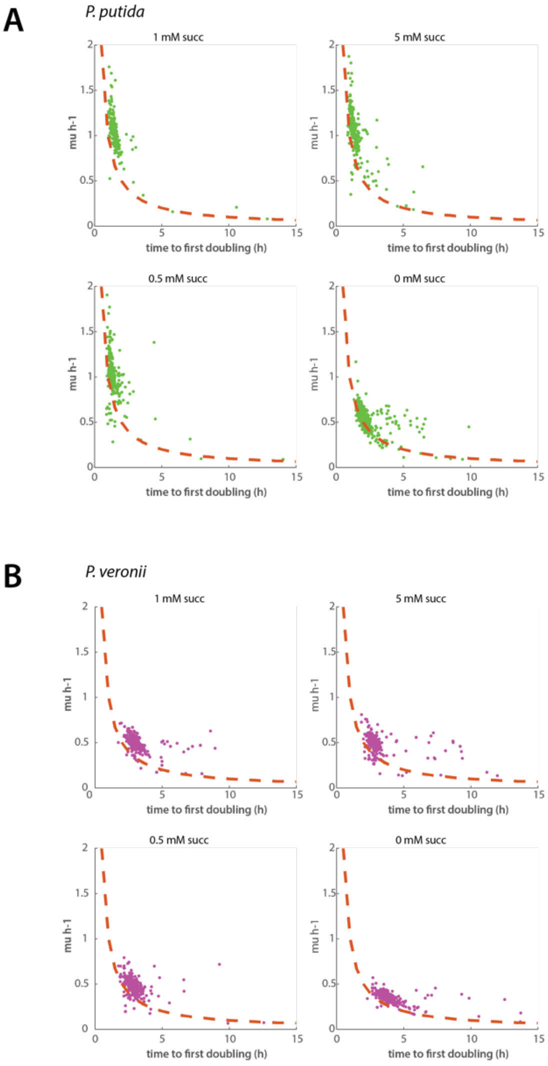


**Supplementary figure 2**. **Relation between the time to first doubling and maximum growth rate, as a function of resource concentration.**

(A) *P. putida* grown on patches with 0, 0.5, 1, and 5 mM succinate. Each green dot indicates a single microcolony measurement. The dashed red curves correspond to the theoretical relation between growth rate and time to first doubling in absence of lag phase. (B) Similar as (A) but for *P. veronii*. Note in both cases a small proportion of founder cells that display a clear lag phase (deviation from the red dashed line).

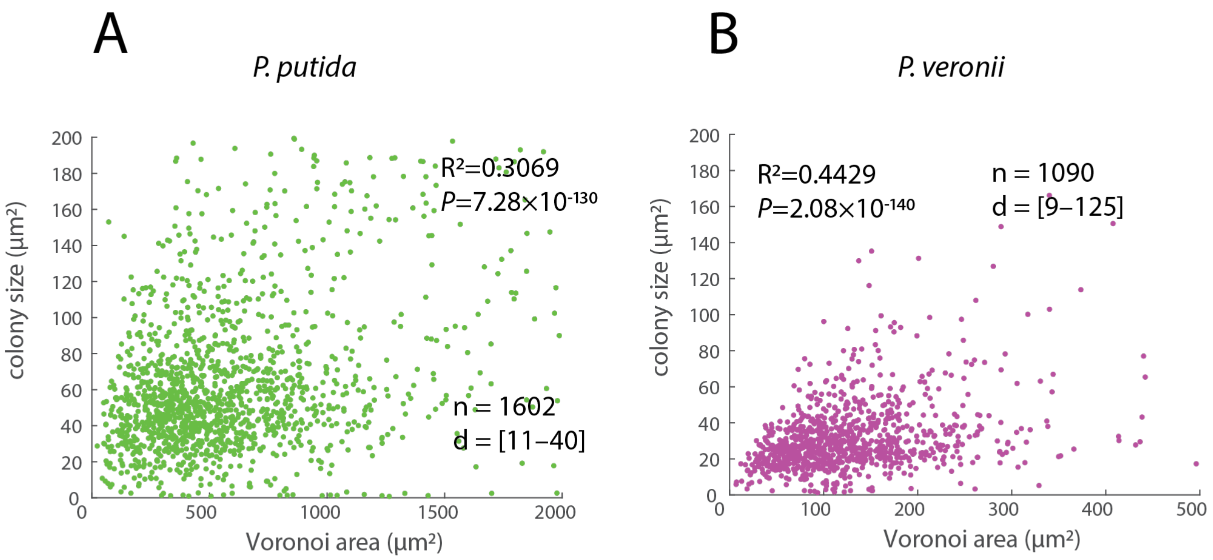


**Supplementary figure 3**. **Correlation between the tesselated Voronoi area at start and the final attained microcolony size in stationary phase.**

(A) *P. putida* grown on patches with 1 mM succinate. Each green dot indicates a single microcolony measurement. n = number of analyzed microcolonies; d = founder cell density range. $R^2$, linear correlation coefficient with corresponding p-value. (B) Similar as (A) but for *P. veronii*.

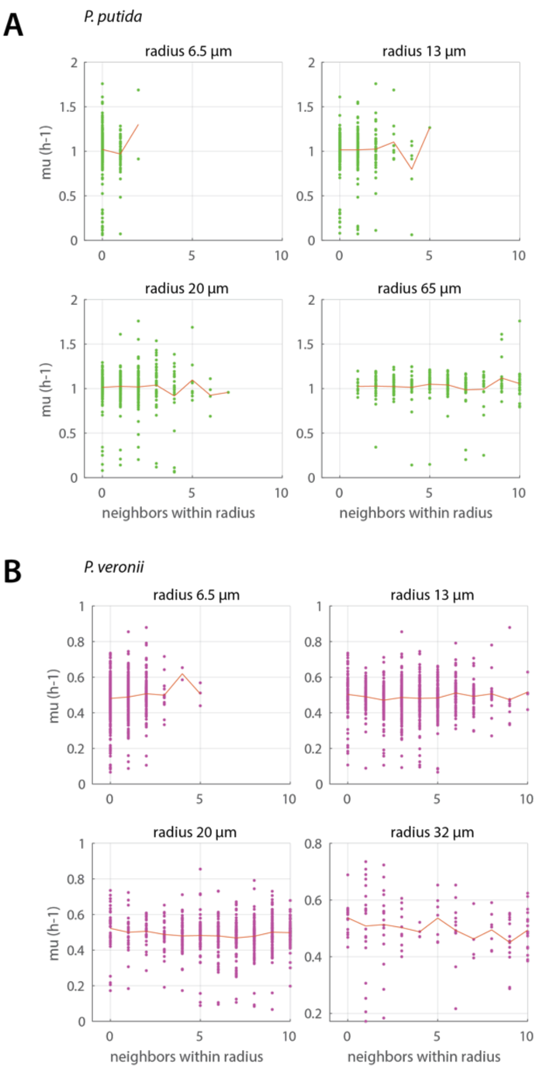


**Supplementary figure 4. Absence of clear impact of the number of neighbours on the maximum growth rates in case of monoculture growth of (A) *P. putida* and (B) *P. veronii*.**

Diagrams display the calculated number of neighbours with the indicated radius around each microcolony founder cell, each dot representing its measured growth rate ($h^{-1}$). The red line combines the mean growth rates as a function of neighbours within the indicated radius.

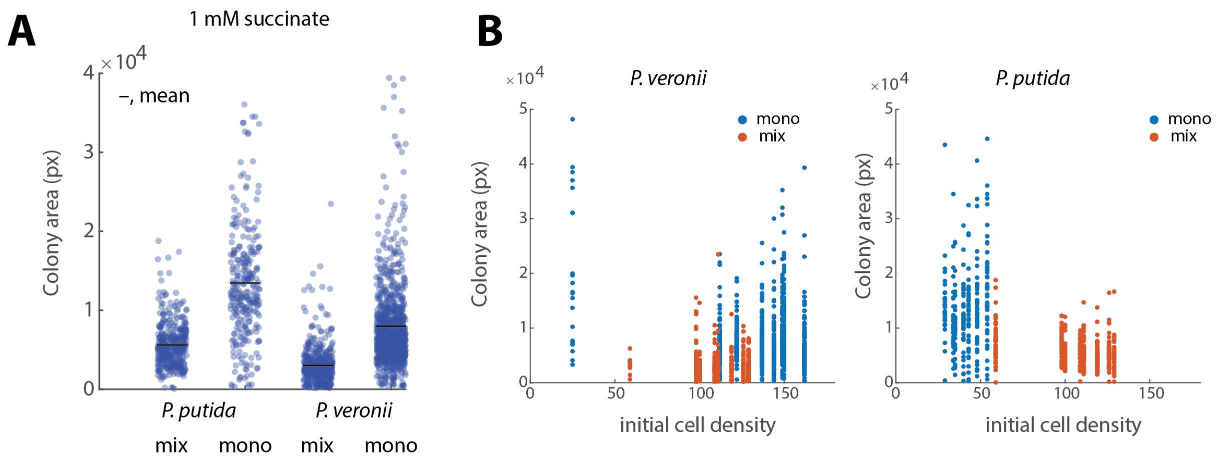


**Supplementary figure 5. Microcolony area differences in mono- versus coculture growth of *P. putida* and *P. veronii* on 1 mM succinate.**

A) Individual measured microcolony areas (dots) in stationary phase from three individual patch experiments, each in with three technical replicates (imaged areas). B) Corresponding initial cell densities and colony area distributions for either *P. veronii* or *P. putida* in mono- or coculture.

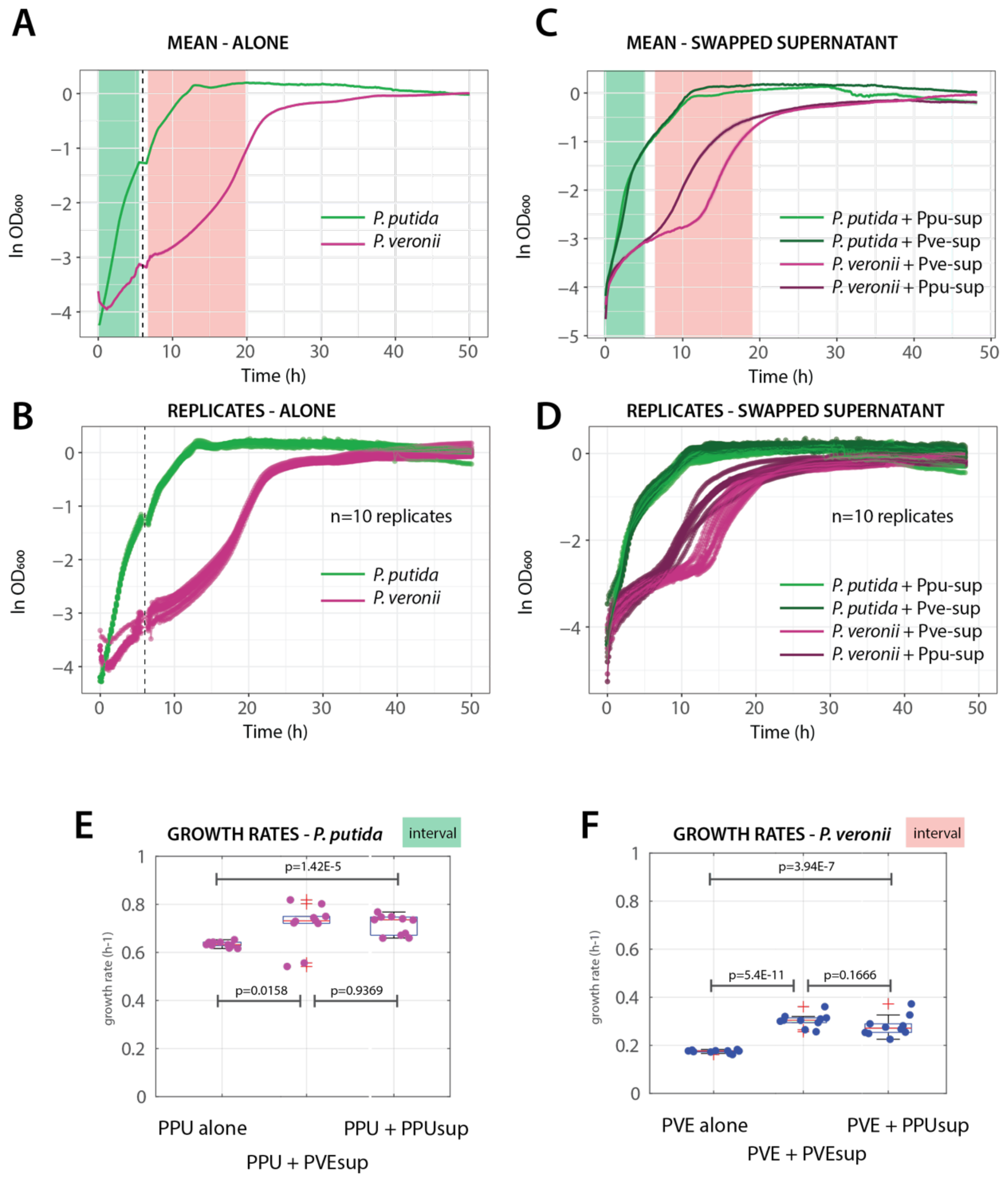


**Supplementary figure 6. Effects of exchanging exponential phase cell-free supernatant on reciprocal growth rates of *P. putida* and *P. veronii*.**

(A) and (B) Mean and individual replicates of 96-well plate reader grown natural logarithm-transformed culture turbidities ($OD_{600}$) *P. putida* or *P. veronii* on 10 mM succinate. Green and red-underlined areas, used for ln-linear regression to find maximum exponential phase growth rates. (C) and (D) Mean and individual replicates of 96-well plate reader grown natural logarithm-transformed culture turbidities ($OD_{600}$) *P. putida* or *P. veronii* on cell-free reciprocal supernatants from cultures grown on 10 mM succinate and harvested at time 6 h (dotted line in panel A). (E) and (F) Derived maximum exponential phase growth rates. P-values from two-tailed t-test (n=10 replicates, each).
